# BOSO: a novel feature selection algorithm for linear regression with high-dimensional data

**DOI:** 10.1101/2020.11.18.388579

**Authors:** Luis V. Valcarcel, Edurne San José-Enériz, Xabier Cendoya, Ángel Rubio, Xabier Agirre, Felipe Prósper, Francisco J. Planes

## Abstract

**Motivation:** With the frenetic growth of high-dimensional datasets in different biomedical domains, there is an urgent need to develop predictive methods able to deal with this complexity. Feature selection is a relevant strategy in machine learning to address this challenge.

**Results:** We introduce a novel feature selection algorithm for linear regression called BOSO (Bilevel Optimization Selector Operator). We conducted a benchmark of BOSO with key algorithms in the literature, finding a superior performance in highdimensional datasets. Proof-of-concept of BOSO for predicting drug sensitivity in cancer is presented. A detailed analysis is carried out for methotrexate, a well-studied drug targeting cancer metabolism.

**Availability:** A Matlab implementation of BOSO is available as a Supplementary Material.

**Supplementary Information:** Supplementary data are available at Bioinformatics online.

## Introduction

High-dimensional datasets are currently an essential part in the biomedical research^1–3^. Much effort has been devoted to developing statistical and machine learning methods able to deal with this complexity^4–8^. Dimensionality reduction and feature selection are the most common strategies to address this issue^9,10^. For linear regression models, we have several approaches to select features. The most popular one is the Lasso regression^11^, which is implemented in different machine learning software packages and integrated in dozens of algorithms for a varied range of biological questions^12–15^.

However, as recently shown in Hastie et al. 2017^16^, the Lasso regression still has substantial room for improvement in high-dimensional datasets. In that work, using synthetic data in a number of conditions, the capacity of several approaches to elucidate the subset of variables that were used to generate the response variable was compared. In particular, they compared Lasso with a recent formulation of the best subset selection approach^17^, which directly addresses the combinatorial problem of identifying the subset of features that more accurately fits the response variable through linear regression. They found that neither approach was significantly better than the other. Interestingly, they concluded that Relaxed Lasso^18^, which combines the solution of Lasso and ordinary linear regression, incorporates the best of both approaches and is, therefore, the most accurate strategy in the literature.

Here, we propose a novel feature selection approach for linear regression called BOSO (Bilevel Optimization Selector Operator). We show that our approach is more accurate than Relaxed Lasso in many cases, particularly in high-dimensional datasets. Proof-of-concept of our approach is applied to predict drug sensitivity in cancer based on RNA-seq data. In particular, a detailed computational and *in-vitro* experimental analysis is presented for methotrexate, a well-studied drug targeting cancer metabolism^19^.

## Results

### The BOSO algorithm

In linear regression, the best subset selection problem addresses the identification of variables correctly related with the response variable. This problem is presented here as a bilevel optimization problem and, for this reason, we call our approach Bilevel Optimization Selector Operator (BOSO). In particular, starting from a total set of *p* features, BOSO searches for the best combination of features of length *K* by solving a bilevel optimization problem, where the outer layer minimizes the validation error and the inner layer uses training data to minimize the loss function of the linear regression approach considered. Here, we chose Ridge regression for the training problem in order to account for multicollinearity in a simpler manner than Lasso; however, the formulation is also presented for ordinary linear regression (see Methods section for details).

In particular, BOSO relies on the observation that the optimal solution of the inner problem can be written as a set of linear equations that depends on the selected features. This observation makes it possible to solve a complex bilevel optimization problem via Mixed-Integer Quadratic Programming (MIQP). This process is repeated for different *K* values until an information criterion, based on the extended Bayesian Information Criterion (eBIC)^20^, is not further improved. eBIC generalizes the Bayesian Information Criterion (BIC) when *p* > *n* (see Methods section), a scenario common in biomedical applications^21^. We adjust here eBIC to take into account the use of Ridge regression instead of ordinary linear regression. Note here that other approaches use validation data to select optimal *K*; instead, BOSO uses validation data to select the best subset of features of length *K* and use information criterion to select the optimal *K*. A conceptual scheme of BOSO for 7 variables can be found in Figure 1.

**Figure 1:**
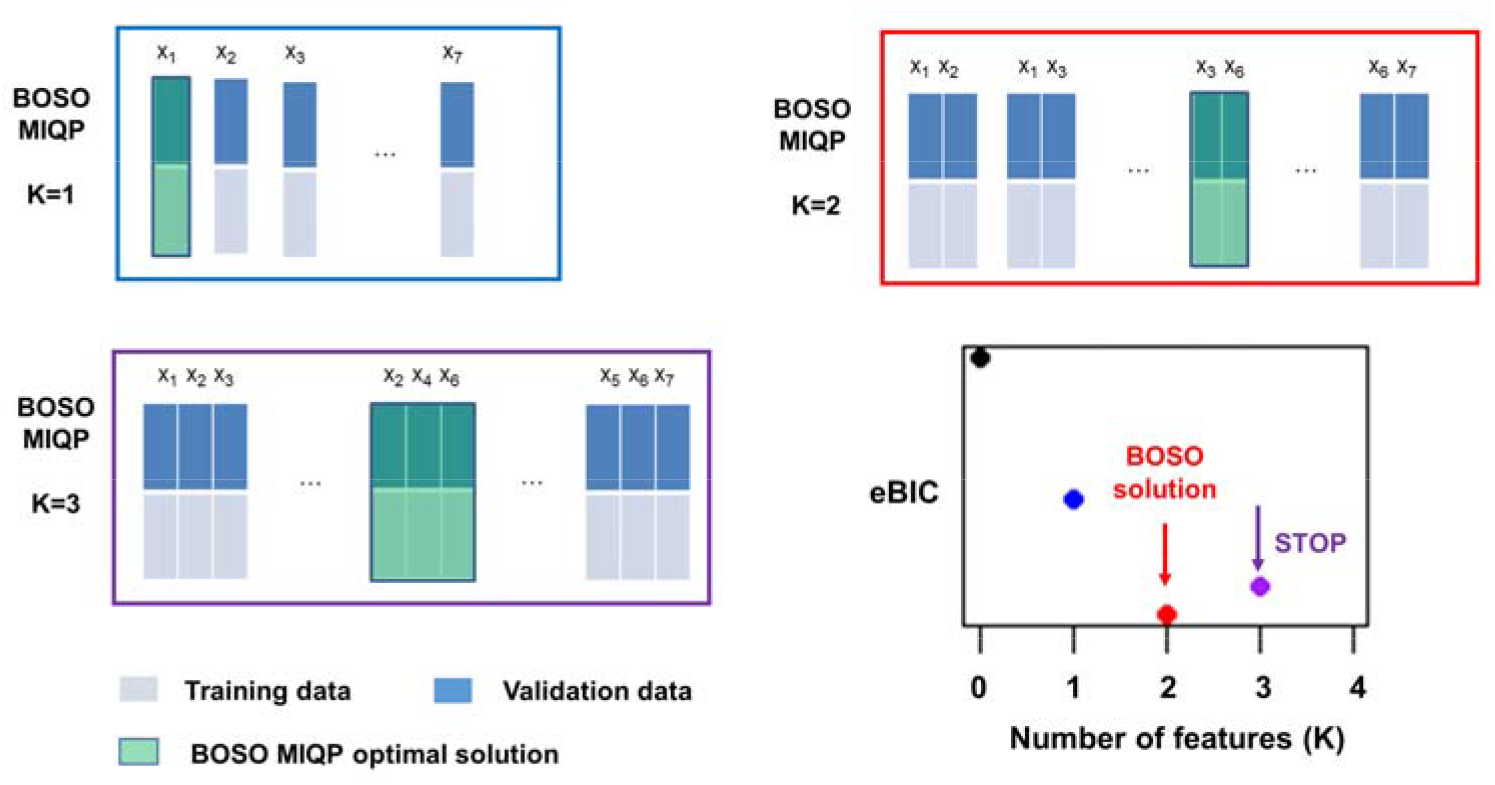
Summary of the BOSO algorithm. An example dataset with 7 features is split into training and validation sets. Green boxes represent optimal selected features for a specific *K* value. For example, for *K*=2, the subset of features that minimizes the validation error is {X_3_, X_6_}. The problem of selecting the best subset of features of length *K* is formulated via mixed-integer quadratic programming (MIQP) (see Methods section) and solved using standard MIQP tools. This process is repeated for each *K* value until our information criterion, which is based on the extended Bayesian Information Criterion (eBIC), is not further improved. Minimal eBIC is found in this example for *K*=2. The final model is derived from Ridge regression with only these two selected variables.

The core MIQP of BOSO addresses a hard combinatorial optimization problem, whose complexity exponentially grows as *p* increases. Current MIQP solvers have been widely developed in the last decade^22^; however, in the case of BOSO, for large problems, they could take long computation times to guarantee optimality. This is also the case of the MIQP approach presented in Bertsimas et al, 2016^17^. Here, we alleviated this issue by iteratively applying BOSO to random blocks of features of length *L* until convergence (see Methods section and Supplementary Fig. 1). With this strategy, we substantially reduced the computation time and managed to apply BOSO to complex problems.

### Benchmarking of feature selection approaches

In order to assess the performance of BOSO, we replicated the same analysis presented in Hastie et al. 2017, where relevant feature selection strategies, including Forward Stepwise^23,24^, Lasso^11^ and Relaxed Lasso^18^, were compared. In that work, they generated synthetic data from a multivariate normal distribution in different settings, which depends on the number of instances, *n*; number of total available features, *p*; actual number of features contributing to the outcome, defined by the sparsity level *s* and their value (beta-type); covariance matrix between features ∑_*ij*_ = *ρ*^|*i*–*j*|^ where *ρ* is the autocorrelation level; and signal-to-noise ratio (SNR level) (see Supplementary Note 1 for further details). In particular, they considered 4 problem settings: low (*n*=100, *p*=10, *s*=5), medium (*n*=500, *p*=100, *s*=5), high-5 (*n*=50, *p*=1000, *s*=5) and high-10 (*n*=100, *p*=1000, *s*=10). These four problem settings were analyzed for different beta-types, autocorrelation level and signal-to-noise ratio.

In particular, we present here the results for one the scenarios considered: beta-type 1, where the *s* contributing features occur at (approximately) equally-spaced indices between 1 and *p* with value 1, the remaining features being equal to 0; and an autocorrelation level between features of 0.35. In this beta-type, actual features contributing to the outcome shows little correlation between them. We tested the same levels of SNR analyzed in Hastie et al. 2017, namely ten values of SNR from 0.05 to 6.00, equally distributed in logarithmic scale. The rest of the cases can be found in Supplementary Figs. 2-21. On the other hand, for this benchmark, we used the expected test error relative to the Bayes error rate, the number of non-zeros estimated coefficients, number of false positives and false negatives, metrics previously used in Hastie et al, 2017 (see Methods section). We also included details as to other evaluation metrics in Supplementary Figs. 2-21.

The comparison of the results obtained with BOSO and with Lasso, Relaxed Lasso and Forward Stepwise clearly demonstrates that BOSO improves their performance as the number of features grows (Figure 2). For the Low setting (*p*=10), BOSO performed the worst in many cases (Figure 2a). However, BOSO reverted this behavior in the Medium setting (*p*=100, Figure 2b), competing with Relaxed Lasso to be the most accurate approach. Importantly, BOSO achieved the best performance in the High-5 setting (p=1000, Figure 2c), obtaining more accurate results than the rest of approaches for all the cases with the exception of SNR=6.00 (Relaxed Lasso was slightly better than BOSO). Finally, a similar behavior is observed in the High-10 setting (p=1000, Figure 2d). According to these results, BOSO is overall more accurate than Forward Stepwise and Lasso and competes with Relaxed Lasso, finding superior accuracy in many cases, particularly in high-dimensional scenarios.

**Figure 2:**
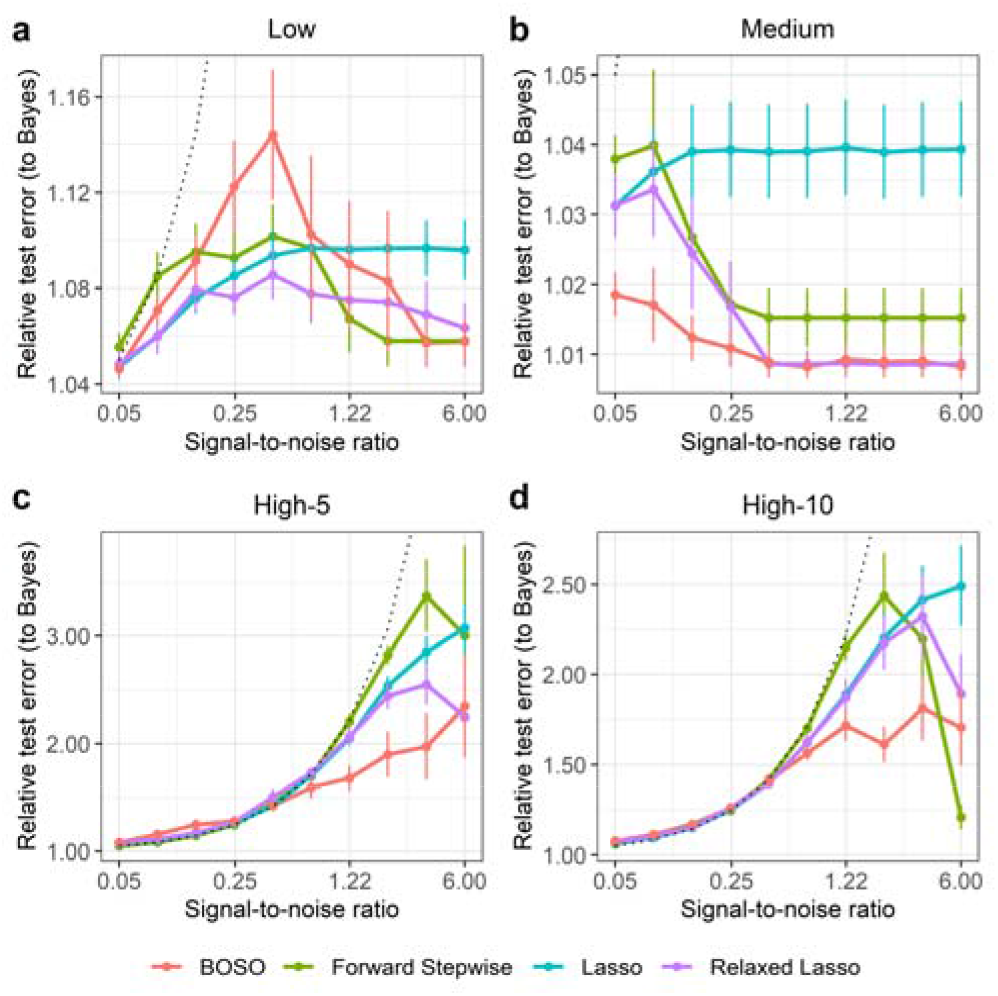
Performance comparison of BOSO with different feature selection algorithms using Relative test error. a) Low setting; b) Medium setting; c) High-5 setting; d) High-10 setting. Dots and bars represent, respectively, the mean and standard deviation of Relative test error across 10 random samples for different SNR values considered. The dotted line is the Relative test error of the null model for each SNR value.

In order to gain insights into the type of model obtained from BOSO, we plotted in Figure 3 the number of non-zeros obtained with each method based on the same simulation as in Figure 2. It can be observed that BOSO generates a more parsimonious model than Relaxed Lasso and Lasso. This is partially derived from our choice of an eBIC-based information criterion to select the size of the model, which is certainly more restrictive than other approaches such as the Akaike Information Criterion (AIC). As a result, BOSO outputs regression models with significantly lower false positives than Lasso and Relaxed Lasso and comparable false negatives (see Figure 4 and Figure 5, respectively). On the other hand, BOSO and Forward Stepwise have similar complexity (Figure 3), but, according to results in Figure 2, Forward Stepwise is less accurate, since it presents higher number of false negatives than BOSO (Figure 5).

**Figure 3:**
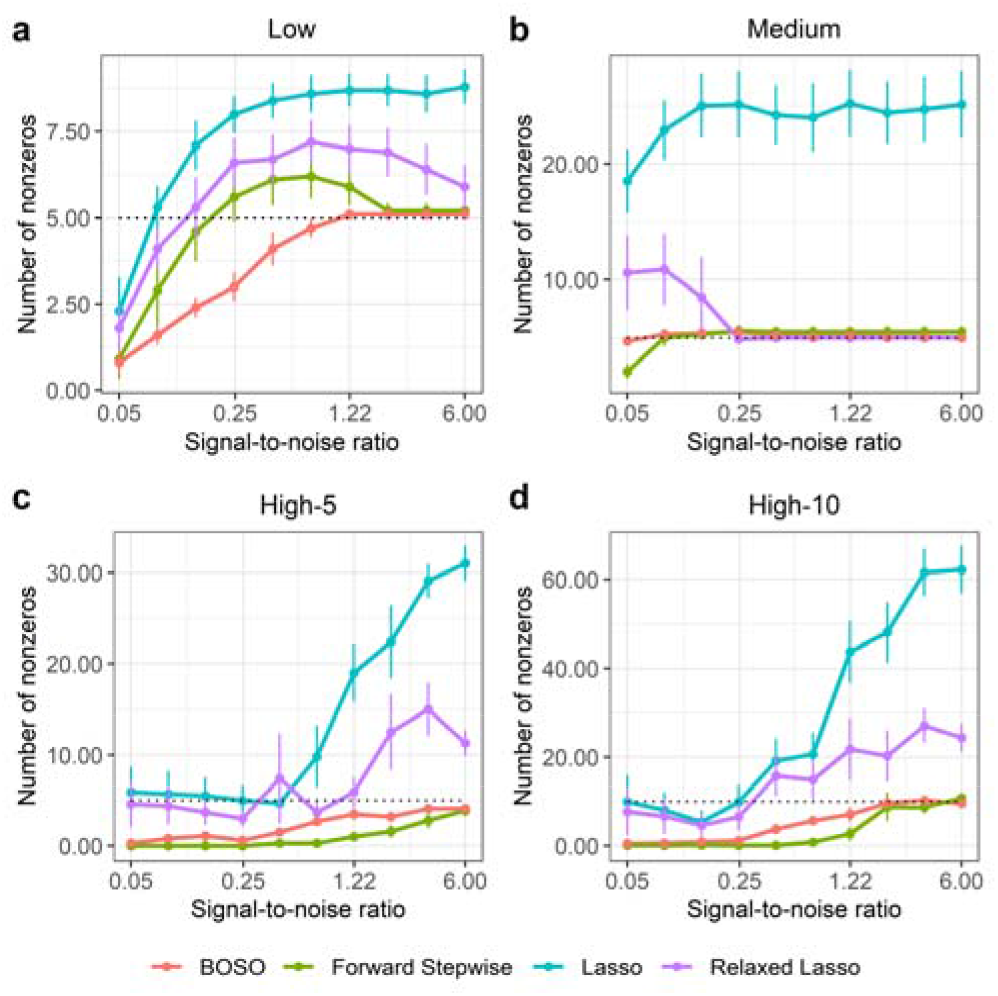
Performance comparison of BOSO with different feature selection algorithms using Number of non-zeros in the 4 problem settings considered. a) Low setting; b) Medium setting; c) High-5 setting; d) High-10 setting. Dots and bars represent, respectively, the mean and standard deviation of Number of non-zeros across 10 random samples for different SNR values considered. The dotted line is the actual value of non-zeros (s) for each SNR value.

**Figure 4:**
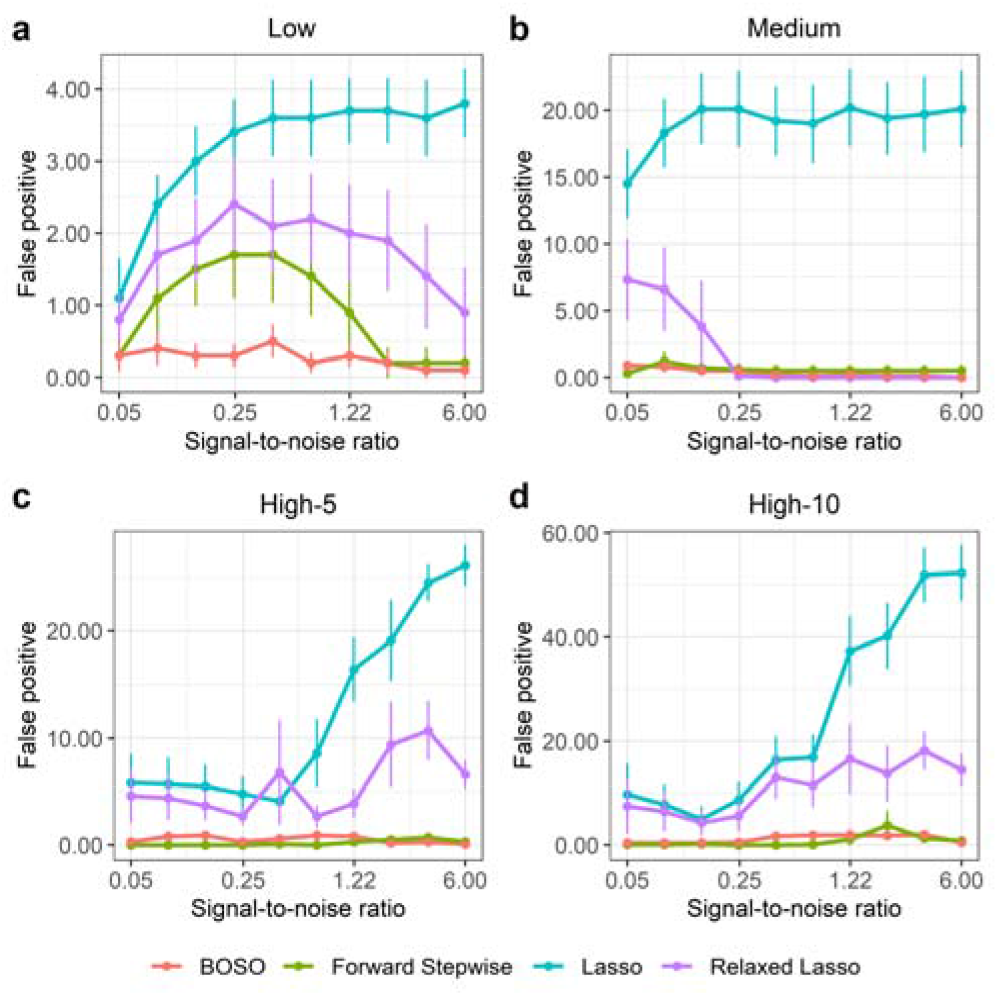
Performance comparison of BOSO with different feature selection algorithms using False Positives in the 4 problem settings considered. a) Low setting; b) Medium setting; c) High-5 setting; d) High-10 setting. Dots and bars represent, respectively, the mean and standard deviation of Number of non-zeros across 10 random samples for different SNR values considered.

**Figure 5:**
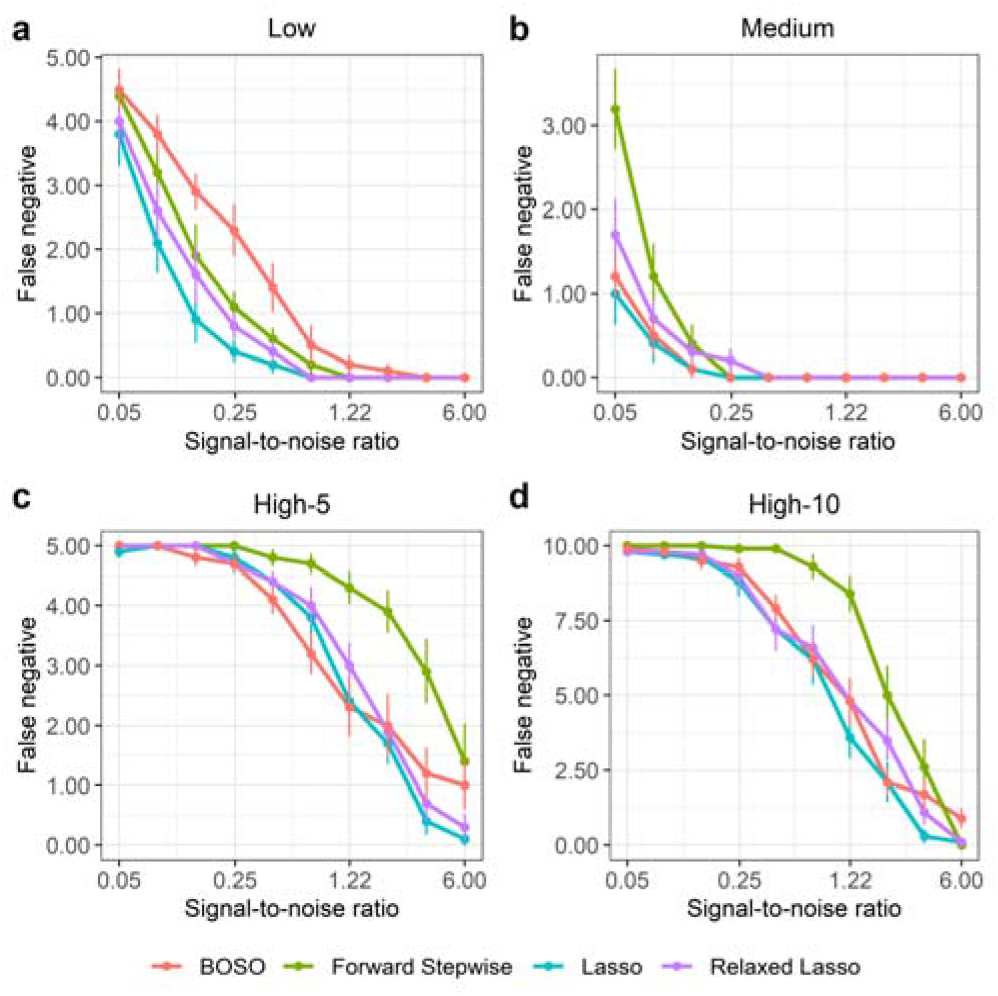
Performance comparison of BOSO with different feature selection algorithms using False Negatives in the 4 problem settings considered. a) Low setting; b) Medium setting; c) High-5 setting; d) High-10 setting. Dots and bars represent, respectively, the mean and standard deviation of Number of non-zeros across 10 random samples for different SNR values considered.

A similar behavior is found for beta-type 2 (see Supplementary Figures 2-21), which defines a more complex situation where actual variables contributing to the outcome are correlated between them. However, we found that BOSO performs worse than Relaxed Lasso for higher correlations in this setting (autocorrelation level 0.70). This is possibly due to the fact that information criterions, such as BIC or eBIC, assumes that variables are independent and they are not prepared for cases where variables present high correlations. This effect is less relevant for more sparse problems, for example, High-5.

With respect to computational effort, even using the random block strategy mentioned above, BOSO requires more time than the rest of the approaches analyzed, particularly Lasso and Relaxed Lasso (Supplementary Table 1). However, BOSO can be run in standard computers, *e.g*. each run in the High-10 setting took us on average 183.501 seconds on a 64□bit Intel(R) Xeon(R) CPU E5-2630 v4 @ 2.20GHz running Linux, setting a maximum of 2 cores and 4 GB of RAM.

In summary, for feature selection: 1) BOSO shows higher sensitivity than Forward Stepwise; 2) BOSO presents higher specificity than Lasso and Relaxed Lasso; 3) BOSO is a computationally feasible approach in large-size problems encountered in biomedical problems.

### BOSO and drug sensitivity in cancer

We applied BOSO to construct a predictive model of Methotrexate (MTX) cytotoxicity in cancer cell lines. To that end, we used 646 cancer cell lines with available IC50 values of MTX from the screenings of the GDSC (Genomics of Drug Sensitivity in Cancer) database^25^ and RNA-seq data from CCLE (Cancer Cell Line Enyclopedia)^26^. After filtering genes with low mean and variance expression (see Methods section), we kept 5304 genes (features) as possible predictors of MTX IC50 (*p*=5304). In order to guide the learning process, cell lines were randomly grouped into training (40%), validation (40%) and test (20%) sets using the R package *caret* (http://topepo.github.io/caret/index.html) for a homogenous distribution of IC50 values. BOSO was applied to training and validation sets and evaluated with test data in 100 different runs. We conducted the same analysis with Forward Stepwise, Lasso and Relaxed Lasso.

It can be observed from Figure 6a that: 1) the models derived from Lasso and Relaxed Lasso have similar correlation in the test data (0.63), slightly superior than BOSO (0.61); 2) Forward Stepwise is the least accurate approach (mean correlation with test data: 0.59). On the other hand, there is a striking difference in the number of features: while BOSO and Forward Stepwise predicted on average 5.65 and 2.34, respectively, Lasso and Relaxed Lasso required more than 50 (Figure 6b). This result is even more extreme when we summarized the features selected (at least once) in the 100 runs: Lasso and Relaxed Lasso selected 1757 and 1448 features, respectively, in at least one of the runs, while BOSO selected 139 features. These results reinforce the conclusions that BOSO generates a more parsimonious model than Lasso and Relaxed Lasso and more accurate model than Forward Stepwise. We repeated the same analysis with 50 drugs available in the GDSC database, finding similar conclusions as the ones obtained for MTX analysis (Supplementary Figure 22).

**Figure 6:**
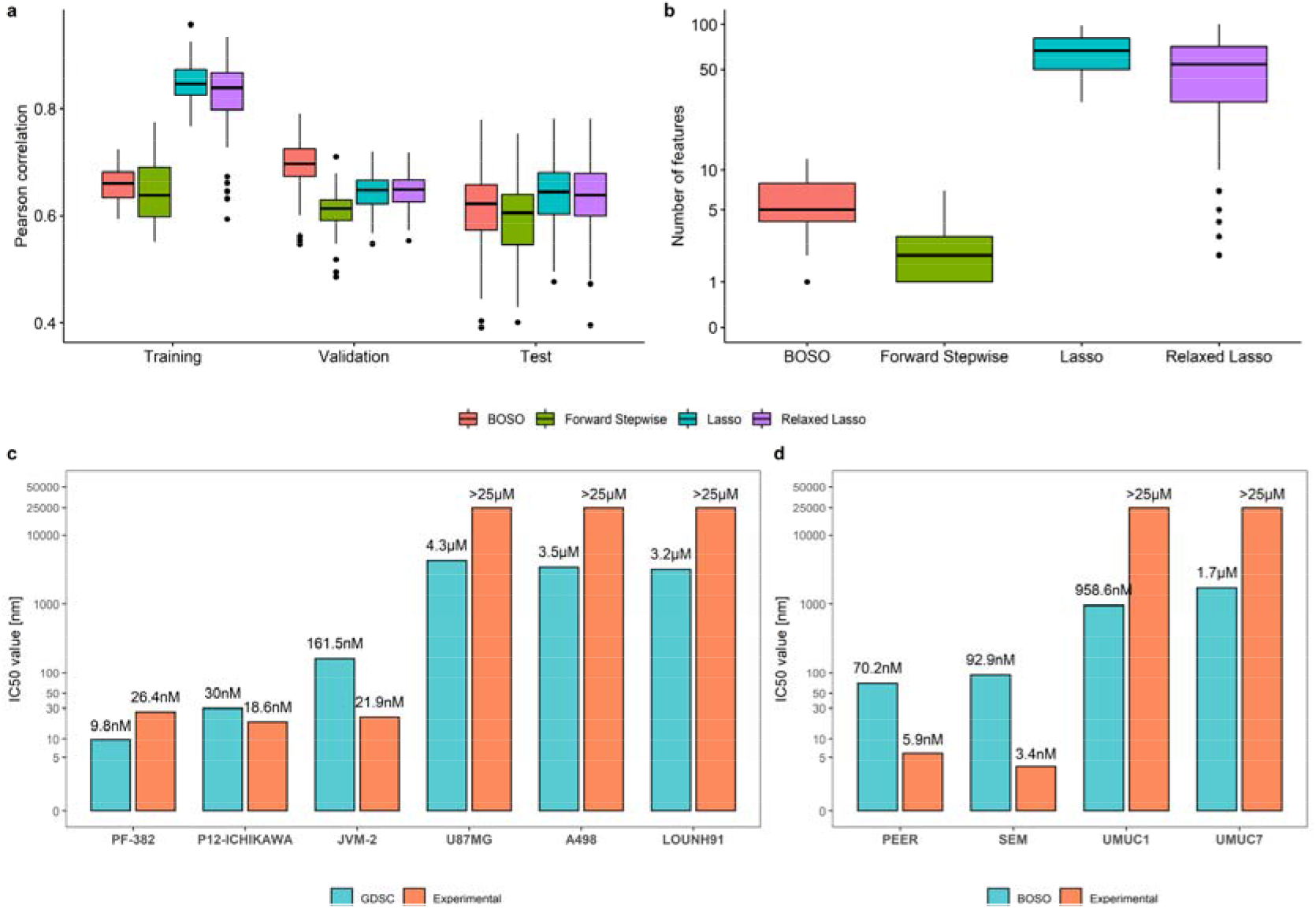
Prediction of Methotrexate cytotoxicity in cancer. Using 100 random partitions of data into training, validation and test sets: a) Pearson correlation obtained with BOSO, Forward Stepwise, Lasso and Relaxed; b) Number of active features selected in the approaches included in Figure 6a; c) Experimental validation of IC50 values provided by the GDSC database for 3 MTX-resistant (U87MG, A498, LOUNH91) and 3 MTX-sensitive (PF-382, P12-ICHIKAWA, JVM-2) cell lines; d) Experimental validation of IC50 values predicted by the BOSO algorithm for 2 MTX-resistant (UMUC1, UMUC7) and 2 MTX-sensitive (PEER, SEM) cell lines, which are not included in the GDSC database.

We exploited the model derived by BOSO to predict MTX IC50 value for 603 cell lines not included in the GDSC database but with RNA-seq data available in CCLE (Supplementary Data). BOSO found clear differences among the distinct cell lines considered, with IC50 values ranging from 10 nM to 923 nM. In addition, we conducted *in vitro* experiments in order to validate our predictive model (see Methods section). First, the IC50 values provided by the GDSC database in 3 MTX-sensitive (PF-382, P12-ICHIKAWA, JVM-2) and 3 MTX-resistant (U87MG, A498, LOUNH91) cell lines (Figure 6c) were validated. This was done because the IC50 values provided by the GDSC database are predicted based on a limited range of experimental screening concentrations^25^. Second, the IC50 values predicted by BOSO in 2 MTX-sensitive (PEER, SEM) and 2 MTX-resistant (UMUC1, UMUC7) cell lines that were not available in the GDSC database (Figure 6d) were assessed in-vitro. A significant agreement between the computational predictions and *in vitro* experiments results were observed, indicating that the linear regression model derived by BOSO can be successfully applied to complete the data provided by the GDSC database.

## Discussion

The feature selection problem is old in machine learning but of high interest nowadays. High-dimensional datasets are proliferating in different domains of science and industry, particularly in biomedical research, where high-throughput −omics technologies, mainly DNA-seq and RNA-seq data, are essential tools for biomarker development in the field of personalized medicine and nutrition. In this context, feature selection approaches are crucial to develop robust machine learning models.

Here, we present BOSO (Bilevel Optimization Selector Operator), a novel feature selection approach for linear regression approaches. BOSO overcomes a complex bilevel optimization problem, linked to the best subset selection problem, based on Mixed-Integer Quadratic Programming. This elegant mathematical transformation is surprisingly novel in the literature. Certainly, existing approaches in the literature addresses the best subset selection using brute force if possible or heuristic methods for more complex problems^27^. Others do not make use of validation data for feature selection and only selects the optimal length, as done in Forward Stepwise. Our strategy is conceptually different and opens new avenues for developing feature selection algorithms in other relevant machine learning tools, such as support vector machines or survival models.

Following the interesting discussion held in the literature^16,17^, BOSO was benchmarked with key feature selection algorithms for linear regression models. BOSO falls between Forward Stepwise and Lasso or Relaxed Lasso. Importantly, BOSO shows higher sensitivity than Forward Stepwise and higher specificity than Lasso and Relaxed Lasso in multidimensional problems, which entails a clear advance in machine learning. This improvement is a mixed result of our proposed MIQP and the choice of our information criterion based on eBIC. However, we think BOSO could be further improved with information criterion taking into account the correlation between the true variables in the model, as currently they are not prepared for this task.

Proof-of-concept of BOSO was accomplished to predict drug sensitivity in cancer. A detailed analysis was presented for methotrexate (MTX), a well-studied drug targeting cancer metabolism. BOSO showed higher accuracy than Forward Stepwise and derived a more parsimonious model than Lasso and Relaxed Lasso, which reinforces again our ability to eliminate false positives. This advantage of BOSO is particularly relevant for biomedical applications, since it simplifies the interpretation, validation and posterior exploitation of results (*e.g*. for the development of combinatorial biomarkers). Finally, we could extend the MTX IC50 values provided by the GDSC database to the remainder 603 CCLE cell lines, providing successful experimental validation for 2 MTX-resistant and 2 MTX-sensitive.

In summary, the results here presented illustrate the value of BOSO for the machine learning community and, in particular, for biomedical research, a field where the number of high-dimensional datasets grows at a frenetic pace. We expect to see the application of BOSO to the great variety of methods where Lasso is being currently applied: predictive models of drug sensitivity, resistance or toxicity, construction of genes regulatory networks, biomarker selection, association studies and other relevant questions.

## Methods

### Bilevel Optimization in Ordinary Linear Regression

Assume a linear regression model with response vector *y* ∈ *R^n^* and design matrix *x* ∈ *R*^*nx*(*p*+1)^, where *p* is the number of predictor variables. The problem of feature selection consists of identifying the subset of predictor variables *Q* that more accurately predicts the response variable *y*. To address this problem with ordinary linear regression, we split the data into training and validation set, namely *y*=[*y^train^, y^val^*] and *X*=[*X^train^, X^val^*], and construct an standard bilevel quadratic optimization model (Eqs. (1)–(4)):

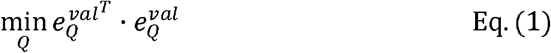

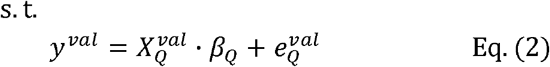

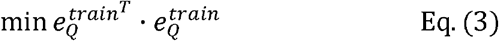

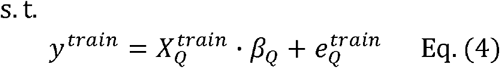

where the inner problem (Eq. (3)–(4)) makes use of the training data for a particular subset of features 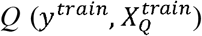 in order to infer its associated optimal parameters *β_Q_* and the outer problem selects the combination of the features *Q* with the lowest validation (generalization) error. Note here that, in bilevel optimization models, the optimal space of the inner problem is a constraint of the outer problem.

The identification of *Q* is a combinatorial problem and approaches in the literature follow a heuristic strategy, such as genetic algorithms^28^. We show below that this bilevel quadratic optimization problem can be reformulated as a mixed-integer quadratic programming model, which can be globally solved with standard optimizers such as IBM ILOG CPLEX. Our approach relies on the observation that the optimal solution of the inner problem can be expressed as a set of linear equations that depends on the selected features. We detail step-by-step this transformation below.

First, let us consider the optimal solutions for the inner problem by assuming that all variables are selected. In that case, following the optimality conditions of ordinary linear regression models (derived from the method of Lagrange multipliers), the inner problem (Eqs. (5)–(6)) can be simplified to a linear set of equations (Eq. (7)):

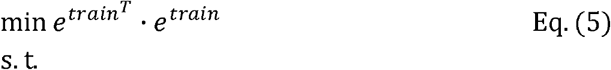

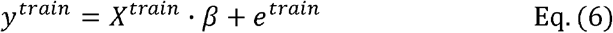

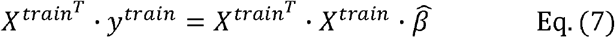

In Eq. (7), we have one equation for each of the features considered plus the intercept (*p*+1 equations). For the sake of simplicity, by making *a* = *X^train^T^^ · y^train^* and *C* = *X^train^T^^·X^train^*, where *a* ∈ *R*^*p*+1^ and *C* ∈ *R*^(*p*+1)*x*(*p*+1)^, we can rewrite the equations algebraically in Eq. (8) and uncoupled in Eq. (9).

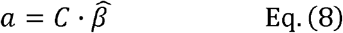

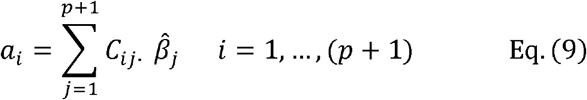

Importantly, coming back to our initial bilevel quadratic optimization problem, the optimality constraints in Eq. (9) only need to be satisfied for the active subset of features *Q* in the inner problem. In other words, if a feature is not considered in the inner problem, then 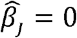 but, additionally, its associated constraint in Eq. (9) must be neglected. These optimality conditions of the inner problem, which depend on the subset of active variables, can be written as a set of linear equations using binary variables *z_i_*, where *z_i_* = 0 if a particular feature *i* is not considered as part of the optimal selection, *z_i_* = 1 otherwise. These equations are written in Eqs. (10)–(13). Note here that *M* is a large positive constant.

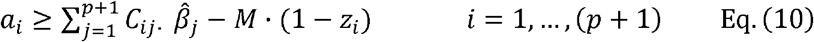

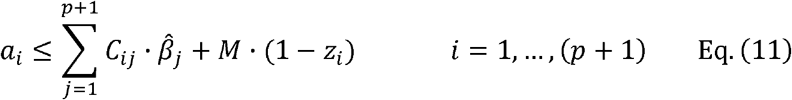

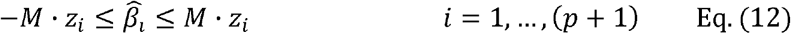

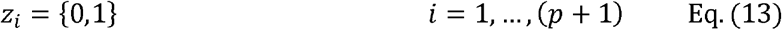

Now we can re-write the bilevel optimization problem as a single mixed-integer quadratic programming problem (MIQP). Our proposed MIQP directly identifies the subset of features that minimizes the validation error given that their associated parameters β are optimal in the training problem. Full details of our MIQP are detailed in Eqs. (14)–(19).

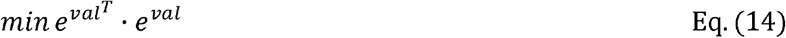

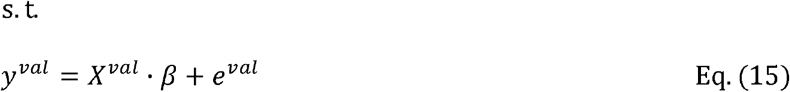

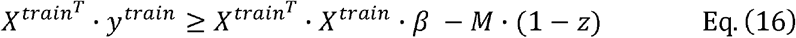

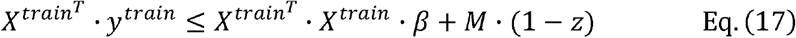

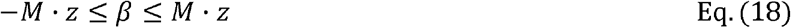

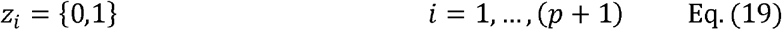

If this MIQP is directly applied, the resulting solution may suffer from overfitting, particularly in cases where the number of features (*p*) is comparable (or higher) to the number of instances (*n*). To avoid this issue, we iteratively apply this MIQP forcing a specific number of features *K* (*K*=1,.., *p*), as shown in Eq. (20), until our information criterion, based on the extended Bayesian Information Criterion of the full model (see below for details), is not further improved.

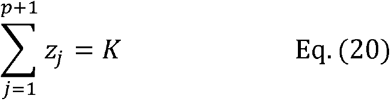

### Bilevel Optimization in Ridge Regression

Similar to ordinary linear regression, the bilevel optimization model associated with Ridge regression is the following:

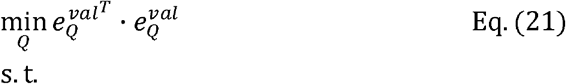

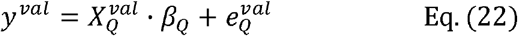

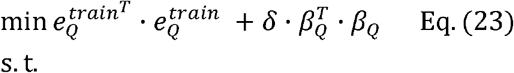

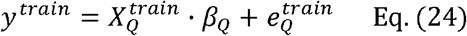

where *δ* is the regularization parameter.

In this case, when all variables are selected, the optimal solution of the inner problem satisfies the following equation (derived from the method of Lagrange multipliers):

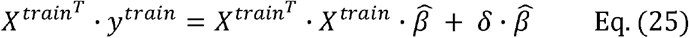

With respect to Eq. (7) in ordinary linear regression, we added the non-linear term 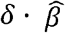. However, for a finite number of *δ* values (*δ*_1_,…, *δ_m_*), as typically used in regularization techniques, we can make it linear through binary variables:

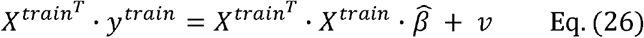

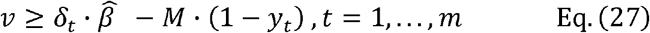

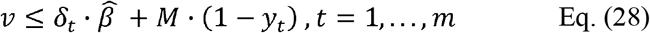

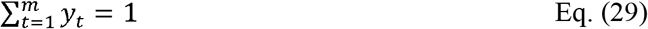

Using *y* variables, we can select the value of *δ* and *v*; in particular, when *y_t_* = 1, then 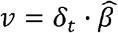; when *y_t_* = 0, the value of *v* is not restricted. As shown in Eq. (29), we can only have one *y* variable as active.

Finally, we can amend Eq. (26) to take into account feature selection. In a similar way as done above for ordinary linear regression, we obtain again a mixed-integer quadratic programming problem that is summarized below:

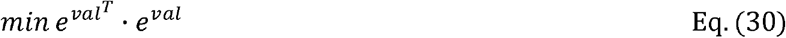

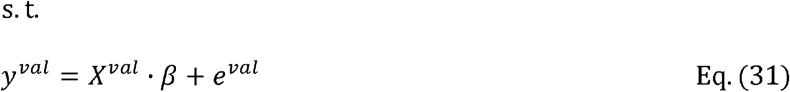

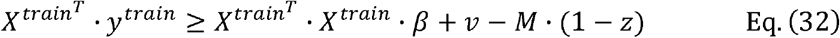

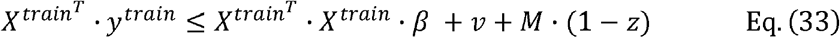

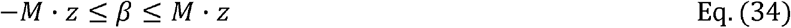

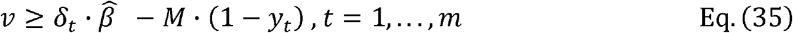

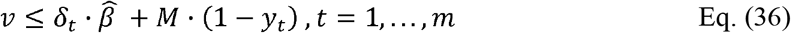

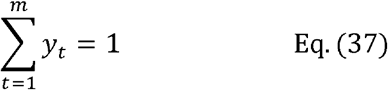

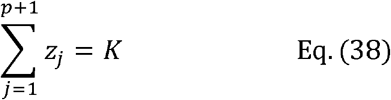

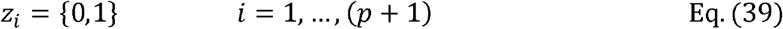

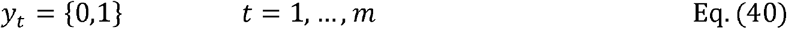

As noted above, we iteratively apply this MIQP, Eqs. (30)–(40), forcing a specific number of features *K* (*K*=1,.., *p*) until our information metric, based on the extended Bayesian Information Criterion (eBIC) of the full model, is not further improved (see next sub-section). With this approach, we obtain the optimal subset of features *Q* and the optimal value of the regularization parameter *δ*. This was the approach used in the Results section. The choice of Ridge regression in the inner layer over ordinary linear regression was done to reduce the variance of the derived model in the event of multicollinearity (high correlation between input variables).

### Extended Bayesian Information Criterion

eBIC is an extension of BIC (Bayesian Information Criterion) for high-dimensional datasets where *p* > *n*. For ordinary linear regression, eBIC is defined in Chen and Chen, 2008^21^, as follows:

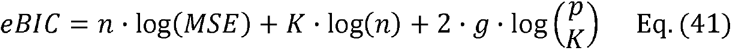

where *n* is the number of instances, *MSE* is the Mean Square Error of the regression model for selected features using both training and validation data, *K* is the number of selected features and *p* is the total number of features. Note here that *g* is a consistency parameter. We used the standard value *g*=0.5 if *p* > *n*; if *p* ≤ *n*, we fixed *g*=0, which is equivalent to Bayesian Information Criterion (BIC).

Here, we modify standard eBIC to consider the use of Ridge regression instead of ordinary linear regression. This was done by substituting the number of features *K* by the effective number of parameters in the model *K_eff_* and degrees of freedom (*df*(*δ*)):

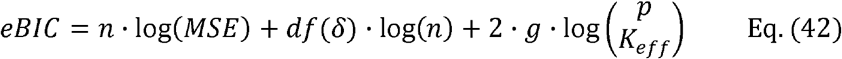

The number of degrees of freedom in Ridge regression is well-known^29^:

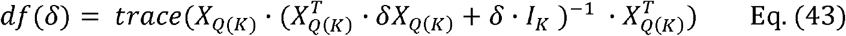

where *X*_*Q*(*K*)_ is the sub-matrix of *X* only including the columns of the *K* features selected. It is easy to observe that if there is no regularization (*δ*=0), the number of effective parameters is precisely *K*. As *df(δ)* will be typically non-integer, we round up *K_eff_* to the nearest integer:

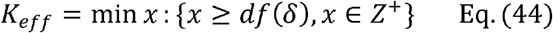

### Computational implementation

In cases with a high number of features, we divide the full set of features into random blocks of features of length *L* (here *L=10*) and apply our MIQP approach described above to each block using *m* different *δ* values (here *m*=10). The selected features in each block are integrated and again divided into random blocks. Our MIQP approach is then applied to each new block. This process is repeated until convergence, namely when the subset of selected features is the same after several iterations or the number of features is less than *L*. In a first stage, in order to select the number of features in each random block, we used BIC instead of eBIC, which is a less restrictive strategy. In a second stage, with the resulting subset of features obtained in the first stage, our random block strategy was repeated using a higher *m* value (*m*=50 for *low* settings, *m*=100 for the rest) and eBIC for feature selection.

We used IBM ILOG CPLEX to solve the MIQP defined by Eqs. (30)–(40). In order to overcome numerical issues derived from the use of the big *M* method in Eqs. (32)–(36), we implemented indicator constraints available in IBM ILOG CPLEX ^30^. MATLAB was used to implement BOSO (see Supplementary Software). We fixed a time limit for each optimization run of 60 seconds on a 64□bit Intel(R) Xeon(R) CPU E5-2630 v4 @ 2.20GHz running Linux, setting a maximum of 2 cores and 4□GB of RAM.

### Accuracy metrics for synthetic data

As defined in Hastie et al. 2017, Relative test error is defined by:

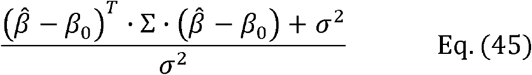

where 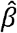 and *β*_0_ represent the estimated and actual value of parameters for the different features considered, respectively; Σ is the covariance matrix of predictor features, whose entry (i,j) is equal *ρ*^(*i–j*)^, being *ρ* the predictor correlation level; *σ*^2^ is the variance of the response variable y. The variance *σ*^2^ is related with the signal-to-noise ratio (SNR) as follows:

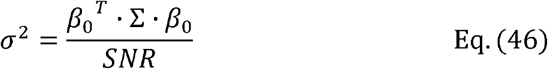

For this comparison in Figure 2, once selected the features and optimal *K* with our MIQP and eBIC, we estimated the coefficients 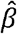 only with training data, as done in Hastie et al. 2017^16^.

### Drug sensitivity in cancer

For the drug sensitivity analysis, RNA-seq data for different CCLE cancer cell lines was downloaded from the DepMap (Dependency Map) portal (www.depmap.org)^31^. Gene expression levels are provided in log2(TPM+1). We kept for further analysis those genes with: 1) mean expression value across the cell lines greater than 1 TPM; 2) variance across the cell lines greater than one unit. IC50 values were also taken from the DepMap portal.

### Cell culture

PF-382, P12-ICHIKAWA, JVM-2, A-498, LOUNH91, U-87MG, PEER, and SEM cell lines were obtained from the DSMZ or the American Type Culture Collection (ATCC) and were authenticated by performing an STR (short tandem repeat) allele profile. UMUC1 and UMUC7 lines were provided by Dr. Paramio at CIEMAT (Centro de Investigaciones Energéticas, Medioambientales y Tecnológicas). U-87MG was cultured with DMEM medium and the rest cell lines were maintained in culture in RPMI 1640 medium supplemented with fetal bovine serum at 37 °C in a humid atmosphere containing 5% CO2. Aside from UMUC1 and UMUC7, the rest of cell lines were tested for mycoplasma (MycoAlert Sample Kit, Cambrex).

### Methotrexate treatment and cell proliferation assay

Methotrexate (S1210) was purchased from Selleckchem (Houston, TX), dissolved in DMSO at 10mM and stored at −80°C.

Cell proliferation was analyzed using the CellTiter 96 Aqueous One Solution Cell Proliferation Assay (Promega, Madison, W). This is a colorimetric method for determining the number of viable cells in proliferation. For the assay, suspension cells were cultured by triplicate at a density of 1×106 cells/mL in 96-well plates (100.000 cells/well, 100μL/well), except for JVM-2 cell line that was cultured at a density of 0.2×106 cells/mL (20.000 cells/well, 100μL/well). Adherent cells were obtained from 80-90% confluent flasks and 100 μL of cells were seeded at a density of 2500 cells /well in 96-well plates by triplicate. Before addition of the compounds, adherent cells were allowed to attach to the bottom of the wells for 12 hours. In all cases, only the 60 inner wells were used to avoid any border effects.

After 96 hours of MTX treatment at different doses, plates with suspension cells were centrifuged at 800 g for 10 minutes and medium was removed. The plates with adherent cells were flicked to remove medium. Then, cells were incubated with 100 μL/well of medium and 20 μL/well of CellTiter 96 Aqueous One Solution reagent. After 1-3 hours of incubation at 37 °C, the plates were incubated for 1-4 hours, depending on the cell line at 37 °C in a humidified, 5 % CO2 atmosphere. The absorbance was recorded at 490 nm using 96-well plate readers until absorbance of control cells without treatment was around 0.8. The background absorbance was measured in wells with only cell line medium and solution reagent. First, the average of the absorbance from the control wells was subtracted from all other absorbance values. Data were calculated as the percentage of total absorbance of treated cells/absorbance of non-treated cells. The GI50 values were determined using non-linear regression plots with the GraphPad Prism v5 software.

## Data availability

The authors declare that all data supporting the findings of this study are available within the article and its Supplementary Materials, or from the corresponding authors upon request.

## Code availability

The code used to generate the results shown in this article can be found in Supplementary Data 2.

## Acknowledgements

This work was supported by the Minister of Economy and Competitiveness of Spain [BIO2016-77998-R], Instituto de Salud Carlos III (ISCIII) [PI16/02024, PI17/00701], CIBERONC (Co-financed with European Union FEDER funds) [CB16/12/00489], ERANET program ERAPerMed [MEET-AML], MINECO Explora [RTHALMY], Cancer Research UK and AECC under the Accelerator Award Programme [C355/A26819] and Fundación Ramon Areces [PREMMAN]. L.V.V. was supported by PFIS [FI17/00297] award from Instituto de Salud Carlos III (ISCIII). X.C. was supported by Basque Government predoctoral grant [PRE_2018.2.0297].

## Author contributions

X.A., F.P. and F.J.P. conceived this study. A.R., L.V.V. and F.J.P. developed BOSO and X.C. and L.V.V. carried out its computational implementation. E.S.J. and X.A. performed the experiments. All authors wrote, read and approved the manuscript.

